# Trajectory-guided dimensionality reduction for multi-sample single-cell RNA-seq data reveals biologically relevant sample-level heterogeneity

**DOI:** 10.1101/2024.09.14.613024

**Authors:** Haotian Zhuang, Xin Gai, Anru R Zhang, Wenpin Hou, Zhicheng Ji, Pixu Shi

**Affiliations:** Department of Biostatistics and Bioinformatics, Duke University; Department of Statistical Science, Duke University; Department of Computer Science, Duke University; Department of Biostatistics, Columbia University

## Abstract

The analysis of single-cell RNA-sequencing (scRNA-seq) data with multiple biological samples remains a pressing challenge. We present `MUSTARD`, a trajectory-guided dimension reduction method for multi-sample multi-condition scRNA-seq data. This all-in-one decomposition reveals major gene expression variation patterns along the trajectory and across multiple samples simultaneously, providing opportunities to discover sample endotypes along with associated genes and gene modules. In data-driven simulation, `MUSTARD` achieves high accuracy in distinguishing sample-level group differences that existing methods fail to capture. `MUSTARD` also demonstrates a robust ability to capture gene markers and pathways associated with phenotypes of interest across multiple real-world case studies.

## Introduction

The emergence of studies that collect multi-sample, multi-condition single-cell RNA-sequencing (scRNA-seq) data provides an unprecedented opportunity to explore the association between phenotype and cell-resolution transcriptomics. For example, several COVID-19 studies collected multiple scRNA-seq data from patients with varied disease severities^1–3^. While several supervised methods have been proposed for cross-condition differential expression analysis^4, 5^, the options for unsupervised analysis of multi-sample scRNA-seq data are still limited. Dimension reduction is commonly used in single-cell analysis^6–9^. Many dimension reduction methods such as t-distributed Stochastic Neighbor Embedding (t-SNE)^10^ and Uniform Manifold Approximation and Projection (UMAP)^11^ have been developed or applied for scRNA-seq data from a single sample, aiming to remove the noise in the original highdimensional gene expression profiles and extract the cellular heterogeneity by inferring a cell-level low-dimensional representation^12^. However, there remains a paucity of methods for conducting dimension reduction on multi-sample scRNA-seq data, accompanied by several notable limitations. First, most existing methods aim to integrate multi-sample data^7,13, 14^, rather than identifying the driving factors that differentiate sample phenotypes. Second, nearly all methods can only infer low-dimensional representation of cells, which can be difficult to connect to sample-level phenotypes. More importantly, current methods do not account for the pseudotemporal information within multi-sample data. Pseudotime analysis has been widely used to study the dynamics of biological processes by ordering cells along a pseudotemporal trajectory^15–17^. A recent study demonstrated that genes exhibit multiple types of changes in a pseudotemporal trajectory across conditions and developed a regression framework for differential multi-sample pseudotime analysis^18^. Therefore, leveraging such pseudotemporal trajectories opens up the possibility of performing dimension reduction for multi-sample scRNA-seq data, enabling the connection between cell-level transcriptomics, sample heterogeneity and phenotypes.

To this end, we present MUlti-Sample Trajectory-Assisted Reduction of Dimensions (`MUSTARD`), a trajectory-guided method for the dimension reduction of multi-sample scRNA-seq data. `MUSTARD` is a pioneering method that utilizes single-cell resolution information to provide unsupervised low-dimensional representation of samples while simultaneously connecting the sample-level heterogeneity with gene modules and pseudotemporal patterns. `MUSTARD` requires three inputs: a gene expression matrix for all cells, a categorical vector indicating which sample each cell belongs to, and the pseudotime values for each cell constructed based on the multi-sample scRNA-seq data using any method appointed by the user according to their cell types of interest. After standard preprocessing steps (e.g., feature selection and scaling), we format the data into an order-3 temporal tensor with sample, gene, and pseudotime as its three dimensions. The tensor is then decomposed into the summation of low-dimensional components, where each consists of a sample loading vector, a gene loading vector, and a temporal loading function (Figure 1a). This all-in-one decomposition is capable of simultaneously revealing sample heterogeneity, major gene expression variation patterns and gene pathway enrichment along the trajectory, thus facilitating sample-level analysis of single-cell data and providing opportunities to discover sample endotypes and corresponding genetic signatures.

**Figure 1.**
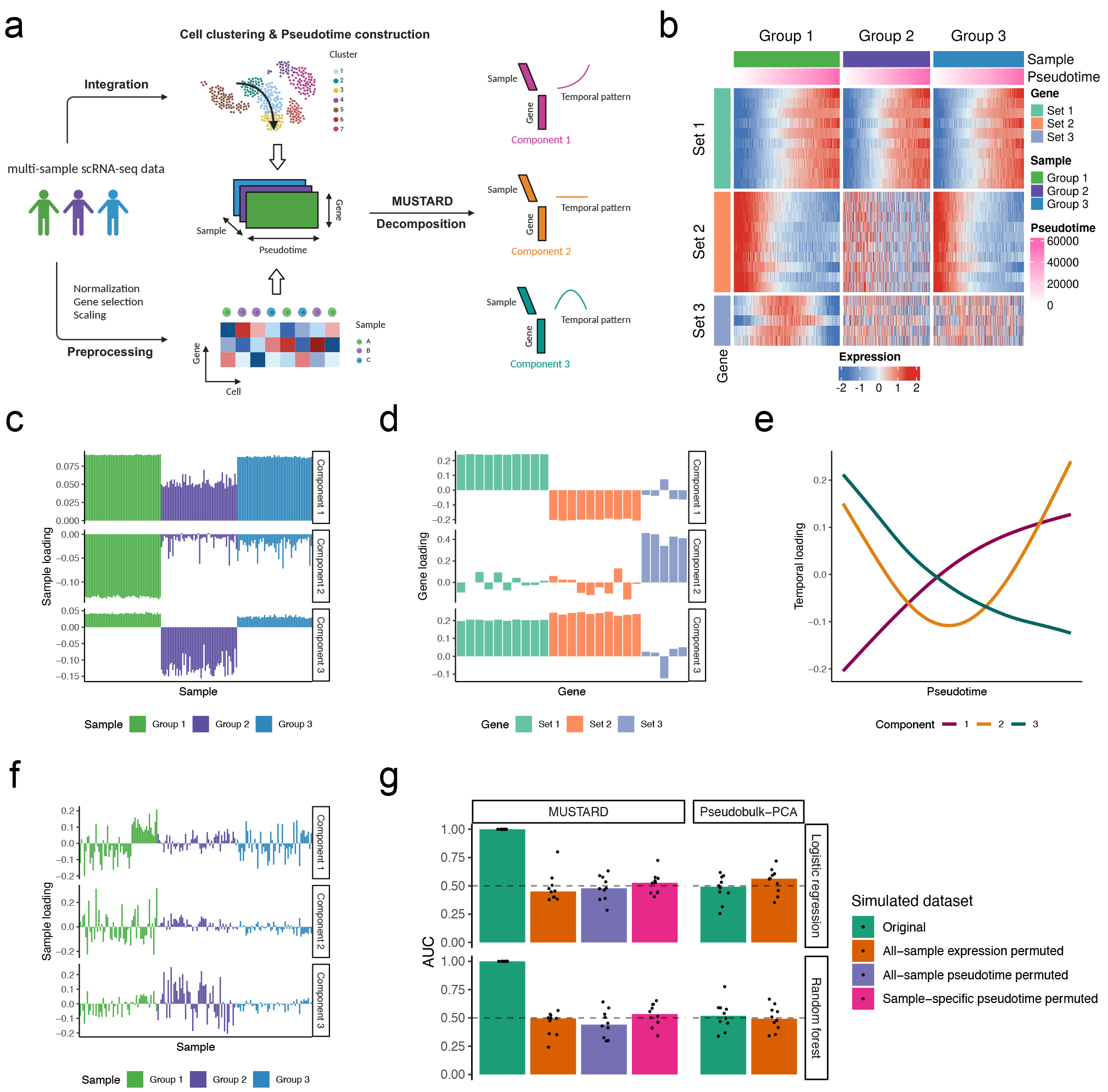
Overview of `MUSTARD` with simulation studies. **a**, A schematic view of `MUSTARD`. **b**, Heatmap showing the multi-sample pseudotemporal expression patterns in the simulated dataset. **c-e**, Sample loadings (**c**), gene loadings (**d**), and temporal loadings (**e**) of the top three `MUSTARD` components from the simulated dataset. The first component captures the monotone trend of genes in Sets 1 and 2, with negative gene loading for the decreasing trend in Set 2, and smaller sample loading for Group 2 due to no trend is in Set 2. The second component captures the U-shaped trend of genes in Set 3 from Group 1. The third component supplements the first component. **f**, Sample loadings of the top three Pseudobulk-PCA components from the simulated dataset. **g**, Performance evaluation of `MUSTARD` and Pseudobulk-PCA using the simulated dataset and three permuted datasets (for details see Methods). Note that when using the Pseudobulk-PCA method, the all-sample pseudotime permuted and sample-specific pseudotime permuted datasets are equivalent to the original dataset.

## Results

First, we demonstrate the capability of `MUSTARD` through a data-driven simulation based on the COVID-Su dataset^1^. This dataset was chosen because a previous study^18^ has constructed a pseudotemporal trajectory from naive T cells to CD8+ T cells and identified genes with significant expression trends along the trajectory. We focused on genes with three different pseudotemporal patterns (Sets 1-3), randomly divided samples into three groups (Groups 1-3), and randomly shuffled the expression of certain genes within each sample in certain groups to differentiate the sample groups (Figure 1b). `MUSTARD` decomposes the simulated multi-sample pseudotemporal data into informative components that depict the differences between gene sets (Figure 1c) and between sample groups (Figure 1d) in terms of their pseudotemporal trends (Figure 1e). Since there is no existing method for sample-level dimension reduction based on scRNA-seq data, we compared `MUSTARD` with what researchers typically do to extract sample-level heterogeneity, i.e., PCA on the aggregated expression across all cells for each sample, referred to as the Pseudobulk-PCA method. Without the cell resolution information, the Pseudobulk-PCA method is incapable of separating any of the groups (Figure 1f).

To further demonstrate the superiority of `MUSTARD` over Pseudobulk-PCA in their ability to capture sample heterogeneity, sample loadings from `MUSTARD` and Pseudobulk-PCA were trained using logistic regression and random forest to predict the phenotypes of unseen samples through 10-fold crossvalidation. The area under the ROC curve (AUC-ROC) was used to evaluate the classification accuracy of both methods. Aside from the original dataset used to assess the power of the methods, we also evaluated their control of type-I error using three types of permuted datasets: all-sample expression permuted, all-sample pseudotime permuted, and sample-specific pseudotime permuted (for details see Methods). Both `MUSTARD` and Pseudobulk-PCA control the type I error rate, i.e., AUC is close to 0.5, in the permuted datasets. However, `MUSTARD` consistently outperforms Pseudobulk-PCA in out-of-sample predictions using the original dataset, achieving a perfect AUC (Figure 1g). The Pseudobulk-PCA method fails to capture the sample-level group differences because it relies on the average expression levels of each sample, which are similar in this simulated dataset. Notably, `MUSTARD` has AUC around 0.5 when only pseudotime was permuted within each sample while sample labels were preserved. This result highlights the importance of utilizing pseudotime in capturing sample-level differences and explains why `MUSTARD` outperforms the Pseudobulk-PCA method. We also evaluated the performance of `MUSTARD` at different noise levels of gene expression (for details see Methods), resulting in a gradual decrease in AUC from 1 to 0.6 (Figure S1).

Next, we applied `MUSTARD` to the original COVID-Su dataset^1^, which sequenced the peripheral blood mononuclear cells (PBMCs) from COVID-19 patients with varying symptom severity (28 mild, 27 moderate, and 19 severe) and 15 healthy donors. Since CD8+ T cell activation is a crucial process of the immune response in COVID-19 patients, we constructed a pseudotemporal trajectory from naive T cells to CD8+ T cells using `TSCAN`^16^, and leveraged them to guide the dimension reduction by `MUSTARD`. The top five `MUSTARD` components capture major temporal patterns including monotone, flat, and single peak (Figure 2a). Component 2 and 3 jointly separate samples from different severity levels (*p*-values = 2.04 *×*10^−3^, 1.78 *×* 10^−5^, respectively, Figure 2b).

**Figure 2.**
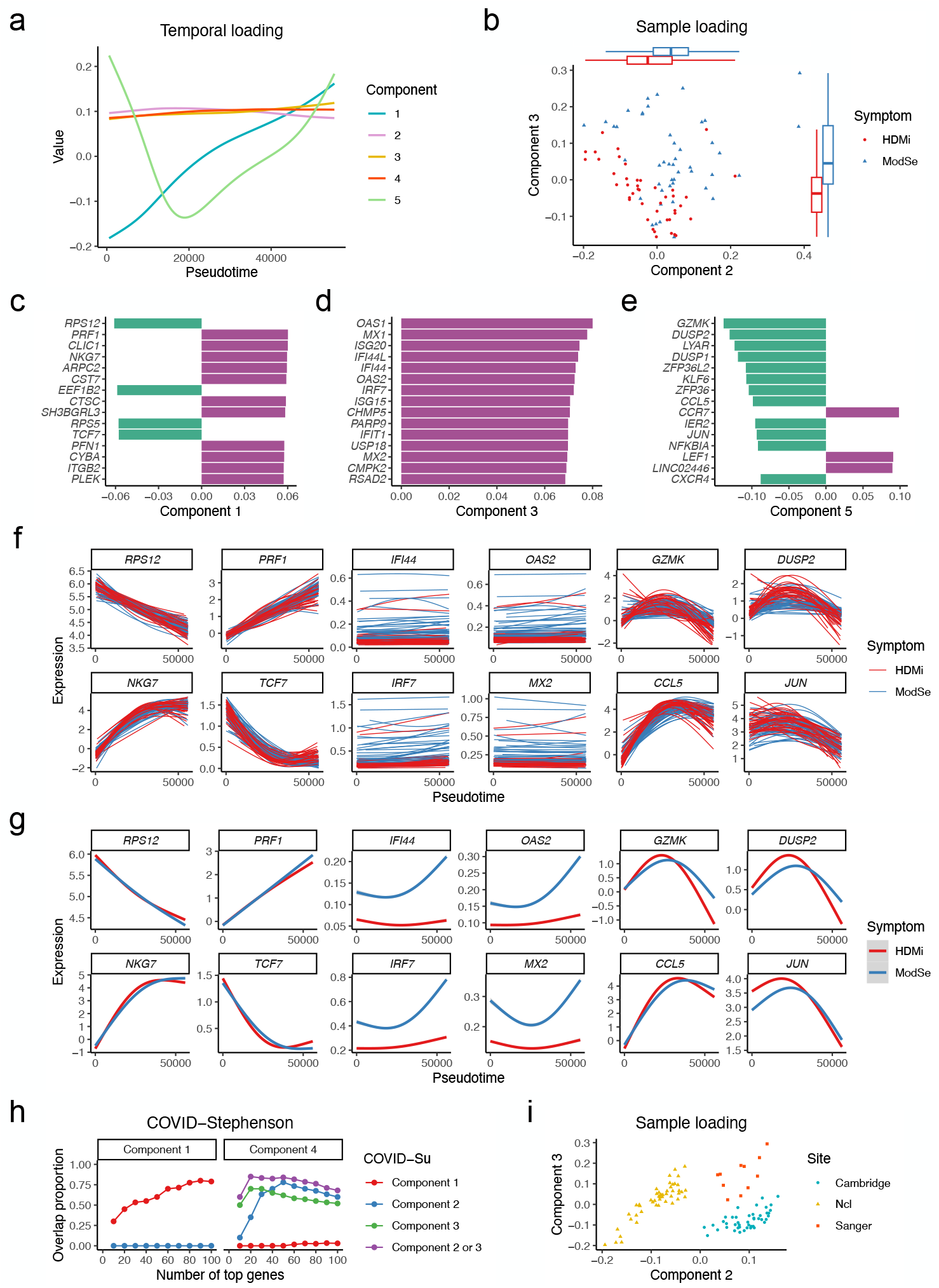
Analysis of COVID-19 studies. **a-g**, Results of the COVID-Su study. **a**, Temporal loadings capture major temporal patterns. **b**, Component 2 and 3 jointly separate samples from different severity levels. *p*-values obtained by Wilcoxon rank-sum test are 2.04*×* 10^−3^ and 1.78 *×*10^−5^, respectively. **c-e**, Top genes in Component 1 (**c**), Component 3 (**d**), and Component 5 (**e**). **f-g**, Example genes’ temporal patterns for each sample (**f**) and each group (**g**). **h**, Proportion of overlapping genes among top *L* genes between the two COVID-19 studies for a range of *L* (for details see Methods). **i**, In the COVID-Stephenson study, Component 2 separates samples from Ncl, and Component 3 further separates samples from Cambridge and Sanger.

We further examine the top genes (ranked by the absolute value of gene loadings) in the aforementioned components. Component 1 is led by genes monotone to pseudotime (Figure 2c). Known genes associated with T cell activation, such as *NKG7* and *TCF7*, ranked at the top due to their increasing and decreasing trend along the trajectory, respectively (Figure 2f-g). Components 2 and 3 are both led by genes differentiating patients with different severity levels (Figure 2d). For example, four genes (*IFI44, OAS2, IRF7, MX2*), ranking top in both Component 2 and 3, are upregulated in moderate or severe patients compared to healthy donors or mild patients along the trajectory (Figure 2f-g), which is consistent with the previous findings indicating CD8+ T cells in moderate patients are programmed to be more terminally differentiated^1, 19^. Component 5 is led by genes with a single-peak temporal pattern, including *GZMK* and *JUN* (Figure 2e-g), which are involved in the transition of cell fate from effector memory T cells to the terminal effector T cell stage^1, 18^.

To demonstrate the reproducibility of `MUSTARD` in real data applications, `MUSTARD` was applied to another independent COVID-19 dataset, referred to as the COVID-Stephenson dataset^2^, which profiled the PBMCs of 110 individuals (24 healthy, 9 mild, 53 moderate, and 24 severe) from three centers including Ncl, Cambridge, and Sanger. Since `MUSTARD` is not supervised by any sample-level variables, there is no guaranteed connection between a single component to the phenotype of interest. Consistent with the COVID-Su study, Component 1 captures monotone trend of genes along pseudotime (Figure S2a). A closer examination of the leading genes in Component 1 of each study reveals a 75% overlap among their top 100 genes (Figure 2c, 2h, S2c). Component 4 separates samples from different severity levels (*p*-value = 2.14 *×* 10^−5^), consistent with Components 2 and 3 of the COVID-Su dataset (Figure S2b). These components have an overlap of up to 80% among their top 20-100 genes, demonstrating the reproducibility of `MUSTARD` on the same biological process across different studies (Figure 2d, 2h, S2d). In contrast, when checking the overlap of components reflecting different aspects of the dataset as a background reference, e.g., Component 2 of the COVID-Su dataset versus Component 1 of the COVID-Stephenson dataset, the percentage of overlap is close to 0 in the leading genes (Figure 2h).

One notable observation from the `MUSTARD` analysis of the COVID-Stephenson study is the batch effect captured by Components 2 and 3, demonstrating a clear separation of patients from three different centers (Figure 2i). Visualizing a cell-level low-dimensional representation (e.g., UMAP) is commonly used as a preliminary step to determine whether batch effects exist in a dataset. However, such a visualization procedure becomes challenging as the number of cells increases from thousands to millions (Figure S2e). `MUSTARD` captures batch effects and mitigates visualization biases by plotting hundreds of samples instead of millions of cells from large multi-sample scRNA-seq data. Additionally, by investigating the leading genes in these components that exhibit the most divergence across three sites, `MUSTARD` elucidates the underlying factors contributing to the batch effect and offers a strategy for minimizing batch effects during the sequencing step (Figure S2f-h). Although numerous methods have been developed to provide low-dimensional representations of cells, such representations are often dominated by the differences between cell types, rendering them less informative regarding sample-level heterogeneity. `MUSTARD`, as the pioneering approach to sample-level dimension reduction, emerges as a potent tool for uncovering unrecognized sample heterogeneity, such as batch effects and endotypes, thereby opening avenues for groundbreaking scientific discoveries.

In addition, the gene loadings obtained from `MUSTARD` can be used to construct pseudotime-informed gene modules. Compared with supervised methods that identify differentially expressed genes one at a time along the trajectory using a spline-based model^15–18^, `MUSTARD` can use its gene loadings to group genes with similar temporal patterns into modules, effectively aggregating signals from multiple genes in connection with both known and unknown source of heterogeneity, which we demonstrate using the COVID-Su study (Figure 3). Different from previous methods that identify gene modules from the original high-dimensional gene expression profiles^21,21^, `MUSTARD` builds the gene-gene corrections based on the gene loadings of top components (3a-b). We evaluate the enrichment of such modules in gene ontology (GO) terms and found distinct enrichment patterns across modules (Figure 3c). The behavior of each gene module along the trajectory can be illustrated through a metagene constructed by averaging the gene expression in each gene module for each sample (Figure 3d-e). For example, the metagenes in Module 1 and 4 show a decreasing and increasing trend, respectively, while the metagene in Module 2 differentiate the samples from different severity levels.

**Figure 3.**
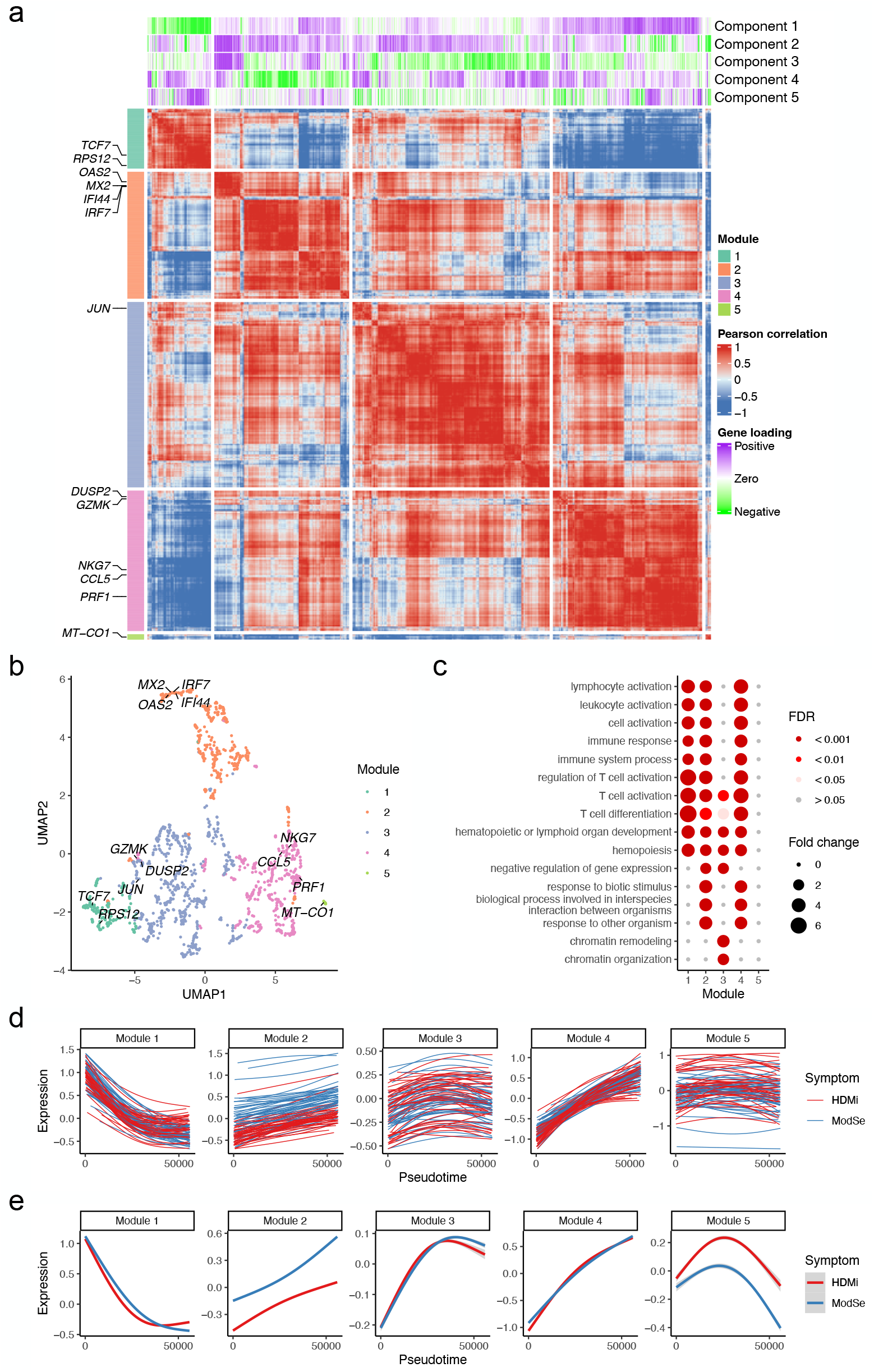
Gene module downstream analysis of COVID-Su study. **a**, Heatmap showing the genegene correlations, determined by gene loadings of the top five components. Example genes in each module are marked. **b**, UMAP showing genes colored by module assignment. **c**, Bubble plot showing the enriched gene ontology (GO) terms identified for each gene module. **d-e**, Metagenes’ temporal patterns for each sample (**d**) and each group (**e**).

In the end, to validate the biological relevance of the genes identified by top `MUSTARD` components, we applied `MUSTARD` to a tuberculosis (TB) dataset obtained from a previous study^22^, where 337,191 memory T cells from 84 males and 100 females were sequenced. Previous studies have found sex to be the dominant grouping factor of subjects, which we will use to validate our findings. To demonstrate the flexibility of `MUSTARD` to use any customized temporal trajectories, instead of inferring pseudotime values using specialized methods, we adopt the trajectory defined in a previous study, which reflects the T cell activation process^18^. Similar to the results from the aforementioned COVID studies, Component 1 captures the monotone temporal pattern (Figure 4a), and the top genes in Component 1 show monotone expression patterns along the trajectory (Figure 4c). Known effector genes, such as *MYO1F* and *ZEB2* increase along the trajectory, while *LTB* and *JUNB* show a decreasing pattern (Figure 4d-e). The sex difference is captured by Component 9 (Figure 4b), whose top 20 genes are all from X and Y chromosomes (Figure 4c-e). In addition, the metagenes constructed from gene module analysis based on the top 10 components reveal several temporal patterns along the trajectory and across samples (Figure 4f-g).

**Figure 4.**
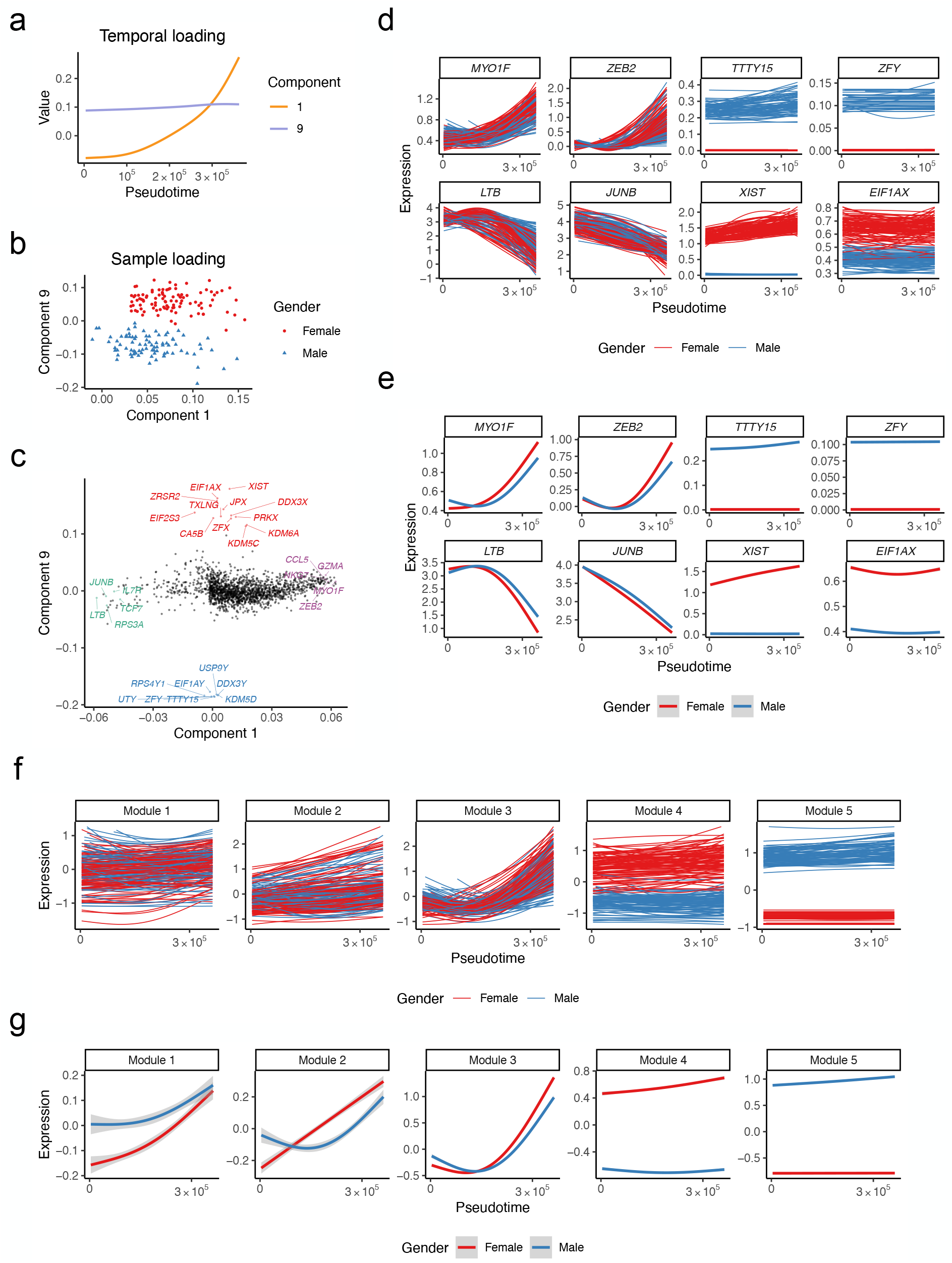
Analysis of TB study. **a-c**, Temporal loadings (**a**), sample loadings (**b**), and gene loadings in Component 1 and 9. The sex difference was captured by Component 9, whose top genes are all from chromosomes X and Y (colored as red and blue, respectively). Top positive (negative) genes in Component 1 are colored as purple (green). **d-e**, Temporal patterns of example genes for each sample and each group (**e**). **f-g**, Temporal patterns of metagenes for modules constructed based on the top 10 components, for each sample (**f**) and each group (**g**).

## Discussion

In summary, `MUSTARD` performs trajectory-guided dimensionality reduction for multi-sample single-cell RNA-seq data. The dimension reduction calculated by `MUSTARD` is dependent on the pseudotemporal trajectories provided by the user. Changing the method for trajectory construction or the starting and ending point of the trajectory may yield different low-dimensional representations of the data. On the other hand, the flexibility of our method to the pseudotime input also allows it to accommodate broader definitions of trajectories subject to the user, such as any meaningful gradient among the cells. `MUSTARD` is designed for utilizing the pseudotemporal information of one trajectory of interest.

When multiple trajectories are considered, the user can apply `MUSTARD` to each trajectory respectively. Alternatively, the user can concatenate the trajectories. An improvement of `MUSTARD` for tree-structured pseudotemporal trajectories will be a focus of our future works.

## Methods

### Data preprocessing and trajectory construction

Before the `MUSTARD` analysis, users need to first integrate multiple scRNA-seq data, followed by cell clustering and pseudotime construction. Users also have the option to define the trajectory with any other one-dimensional numeric values of interest. Prior to the `MUSTARD` analysis, standard preprocessing steps including normalization, identification of highly variable genes (HVGs), and scaling are also required. Since our method uses a tensor decomposition designed for continuous data, data imputation (e.g., `SAVER`^23^) and cell binning are recommended to mitigate the sparsity of the original data. Note that users should use the original gene expression values instead of the corrected expression values from integration methods, which remove the biological variation across samples.

### COVID-Su dataset

The COVID-Su dataset^1^ consisting of 270 samples from 129 COVID-19 patients and 16 healthy donors was downloaded from the ArrayExpress website https://www.ebi.ac.uk/biostudies/arrayexpress/studies/E-MTAB-9357. `SAVER`^23^ (version 1.1.2) was used to impute the library-size-normalized count data from each sample, followed by log2-transformation. Consistent with a previous study^18^, CD8+ T cells were annotated based on the `SAVER`-imputed values of *CD3D* and *CD8A*. Cells with fewer than 500 expressed genes, fewer than 2,000 counts, or more than 10% mitochondrial counts were filtered out. We retained 161 samples with more than 500 cells and 100 CD8+ T cells. `Seurat`^7^ (version 3.2.1) was used to integrate data from multiple samples and `TSCAN`^16^ (version 1.7.0) was applied to construct a pseudotemporal trajectory using 55,953 naive and CD8+ T cells with default settings. The trajectory was divided into 50 consecutive intervals (bins) of equal length because we found that 50 bins provided enough resolution to characterize the pseudotemporal trajectory of the cells in this dataset, and more bins did not lead to meaningful changes in the results. For a given gene and a given sample, the expression value within each bin was represented by the median of the log2-transformed `SAVER`-imputed expression values in cells with pseudotime falling into the bin, and the corresponding pseudotime of the binned cell was represented by the midpoint of the bin. After cell binning, genes with expression values higher than 0.1 in at least 5% of cells were retained. We next applied a gene-specific local polynomial regression (LOESS) to fit the relationship between the standard deviation and the mean of expression values. Genes with positive residuals were selected as HVGs, and then standardized to have zero mean and unit variance. Because a proportion of patients have repeated sampling, we only focused on the 89 samples collected at the first time point from participants, consisting of 28 mild, 27 moderate, and 19 severe COVID-19 patients, and 15 healthy donors. After preprocessing, a final set of 1,600 genes on 3,719 metacells was formatted into an order-3 temporal tensor with 89 samples, 1,600 genes, and 50 pseudotime values as its three modes. Since previous studies have shown a high consistency between healthy to mild samples, and between moderate to severe samples^1^, we dichotomized the severity levels into two groups, i.e., healthy or mild (HDMi) and moderate or severe (ModSe).

### COVID-Stephenson dataset

The COVID-Stephenson dataset^2^ consisting of 143 samples from three centers (Ncl, Cambridge, and Sanger) was downloaded from the ArrayExpress website https://www.ebi.ac.uk/biostudies/arrayexpress/studies/E-MTAB-10026. Similar to the COVID-Su dataset, we only focus on the 110 samples collected at the first time point from 9 mild, 53 moderate and 24 severe COVID-19 patients, and 24 healthy donors. These 110 samples comprised 51 from Ncl, 48 from Cambridge, and 11 from Sanger. The severity of COVID-19 in each sample was standardized based on the World Health Organization (WHO) ordinal score, which was used in the COVID-Su dataset. We followed the same processing steps as described in the COVID-Su dataset, except that the pseudotemporal trajectory was constructed using Harmony^13^-integrated low-dimensional embeddings and cell type annotations obtained from the original data. After feature selection and scaling, a final set of 1,833 genes on 4,614 metacells was formatted into an order-3 temporal tensor with 110 samples, 1,833 genes, and 50 pseudotime values as its three modes.

### Tuberculosis (TB) dataset

The TB dataset^22^ consisting of 500,089 memory T cells from 259 individuals was downloaded from the GEO website https://www.ncbi.nlm.nih.gov/geo/query/acc.cgi?acc=GSE158769. We retained 184 samples from 84 males and 100 females with more than 1,000 cells, using the same filtering criteria as in a recent study^18^. A pseudotemporal trajectory reflecting the T cell activation process was directly obtained from the authors of `Lamian`^18^. After the same processing steps, a final set of 1,790 genes on 9,038 metacells was formatted into an order-3 temporal tensor with 184 samples, 1,790 genes, and 50 pseudotime values as its three modes.

### Simulations

To generate a realistic dataset with multiple samples and different pseudotemporal patterns, we first manually selected a subset of genes with desired pseudotemporal patterns in the COVID-Su dataset. To be specific, we selected 10 genes with increasing trend (Set 1), 10 genes with decreasing trend (Set 2), and 5 genes with a single peak (Set 3) across all samples. The gene expression is fitted along pseudotime by the generalized additive model (GAM). GAM was implemented using the R function `gam()` from the R package `mgcv` (version 1.8.41) with the formula *y* ∼ *s*(*t, k* = 3), same as `TSCAN`^16^. Next, we randomly divided 161 samples into three groups (Groups 1-3). For genes in Set 2, we permuted the expression within each sample in Group 2. For genes in Set 3, we permuted the expression within each sample in Group 2 and Group 3. Finally, a Gaussian noise with zero mean and standard deviation of 0.1 was added back to the fitted expression values. To evaluate the performance of `MUSTARD` at different noise levels, the standard deviation of the Gaussian noise was increased incrementally from 2 to 5. We performed the same binning and scaling steps as described above. The top three `MUSTARD` components were used for downstream analyses.

To compare with the Pseudobulk-PCA method, we took the average expression across all cells for each sample, and then standardized each gene’s expression to have zero mean and unit variance. PCA was then performed to extract the top three principal components. Building upon the simulated dataset above, we then created three permuted datasets (all-sample expression permuted, all-sample pseudotime permuted, and sample-specific pseudotime permuted) that ignore certain information. For all-sample expression permuted dataset, the gene expression of cells was permuted across all samples. For all-sample pseudotime permuted dataset, the pseudotime of cells was permuted across all samples. For sample-specific pseudotime permuted dataset, the pseudotime of cells was permuted within each sample. Note that when using the Pseudobulk-PCA method, the all-sample pseudotime permuted and sample-specific pseudotime permuted datasets are equivalent to the original dataset.

### MUSTARD algorithm

Denote **Y** as the order-3 temporal tensor after the preprocessing steps, with *m* samples and *n* genes, and *y*_*ijt*_ as the expression level of gene *j* from sample *i* at pseudotime *t* ∈ *T*_*i*_ which can be different across samples. The core of the `MUSTARD` algorithm is to decompose **Y** using an approximately CP low-rank structure^24^.

Denote 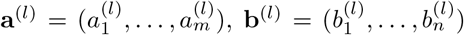, and ***ξ***^(*l*)^ as the sample loading, gene loading, and temporal loading for component *l*, respectively. We estimate these loadings by minimizing the objective function

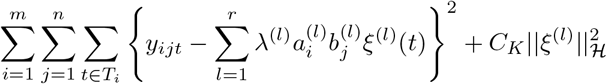

subject to the following constraints:

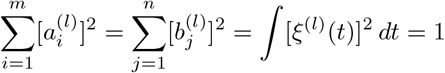

where *r* is the number of low-rank components to approximate **Y**, and ***λ*** = (*λ*^(1)^, …, *λ*^(*r*)^) quantifies the contribution of each component. ||*ξ*|| _ℋ_ is the reproducing kernel Hilbert space (RKHS) norm of function *ξ*(*t*) with the rescaled Bernoulli polynomial kernel

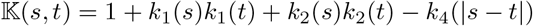

where 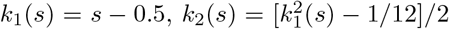, and 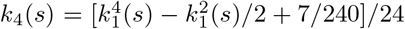. A tuning parameter *C*_*K*_ controls the smoothness of *ξ*(*t*).

For each component *l* = 1, …, *r* sequentially, we perform the following Steps 1 to 3 to estimate **a**^(*l*)^, **b**^(*l*)^, and ***ξ***^(*l*)^.

Step 1: Initialize 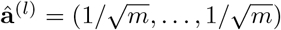. Set 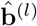 as the first left singular vector of mode-2 matricization of **Y**.

Step 2: Minimize the following function by iteratively updating 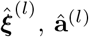, and 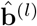 respectively until convergence:

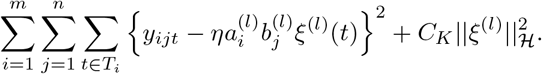

(2a) Update 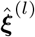 by minimizing

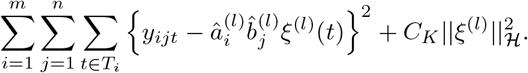

with kernel ridge regression. Then normalize 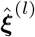 to 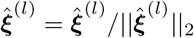.

(2b) Update **â**^(*l*)^ by

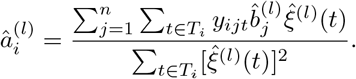

Then normalize **â**^(*l*)^ to **â**^(*l*)^ = **â**^(*l*)^/|| **â**^(*l*)^||_2_.

(2c) Update 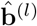 by

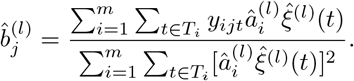

Then normalize 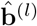 to 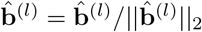.

Step 3: Estimate *η* by minimizing

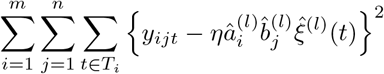

with the least squares method. Then update **Y** by 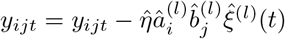

Step 4: Estimate ***λ*** by minimizing

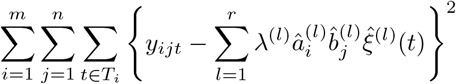

with the least squares method.

## Downstream analyses

### Sample phenotype prediction

Denote **Y**_train_ and **Y**_test_ as the temporal tensor from the training samples and testing samples, respectively, where the trajectory was defined on the entire samples. For each component *l* = 1, …, *r* of **Y**_train_, denote 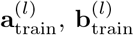, and 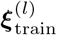 as its sample loading, gene loading, and temporal loading, respectively. Suppose that **Y**_train_ and **Y**_test_ share the same gene loading and temporal loading, we then directly estimate the sample loading of **Y**_test_ (i.e., 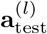) by plugging 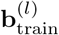 and 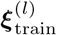 in Step 2 of the `MUSTARD` algorithm. The phenotype of the testing samples will be predicted by applying classification methods such as logistic regression and random forest trained by 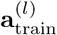 to 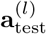.

Logistic regression was implemented using the R function `glm()` from the R package `stats` (version 4.2.2). For multi-group prediction (≥ 3), multinomial logistic regression was implemented using the R function `multinom()` from the R package `nnet` (version 7.3.18). Random forest was implemented using the R package `randomForest` (version 4.7.1). AUC-ROC was calculated using the R package `pROC` (version 1.18.4).

### Gene module identification

Given ***λ*** = (*λ*^(1)^, …, *λ*^(*r*)^) and **b** = (**b**^(1)^, …, **b**^(*r*)^), we denote **B** = (**B**^(1)^, …, **B**^(*r*)^) as the gene embedding matrix, where **B**^(*l*)^ = *λ*^(*l*)^**b**^(*l*)^ and *l* = 1, …, *r*. Users have the option to use any clustering method on the gene embedding matrix with preferred distance measures. In this study, hierarchical clustering with Pearson correlation distance measure was implemented using the R function `hclust(method = “average”)` from the R package `stats` (version 4.2.2). UMAP was implemented using the R function `umap(metric = “correlation”)` from the R package `uwot` (version 0.2.1). GO enrichment was performed using the R package `topGO` (version 2.50.0). The R package `ComplexHeatmap` (version 2.15.4) was used for visualization.

### Cross-study overlap proportion

To quantify the similarity of the top-ranked genes identified in different studies, the overlap proportion in Figure 2h is defined as the proportion of top-ranked genes identified in both studies. Specifically, denote *A* and *B* as the set of top *L* genes identified in two studies. |*A*| = |*B*| = *L*, where |.| is the cardinality of a set. The overlap proportion is defined as 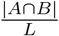. The overlap proportion was calculated for different choices of *L*.

## Availability of data and materials

All datasets used in the study are publicly available. The R package `MUSTARD` with a detailed user manual is publicly available at https://github.com/haotian-zhuang/MUSTARD. The source code to reproduce the results is available at https://github.com/haotian-zhuang/MUSTARD_Paper. BioRender (BioRender.com) was used for generating Figure 1a under a paid subscription, and the publication agreement number is QX26VWQQEY.

---

**Figure S1:**
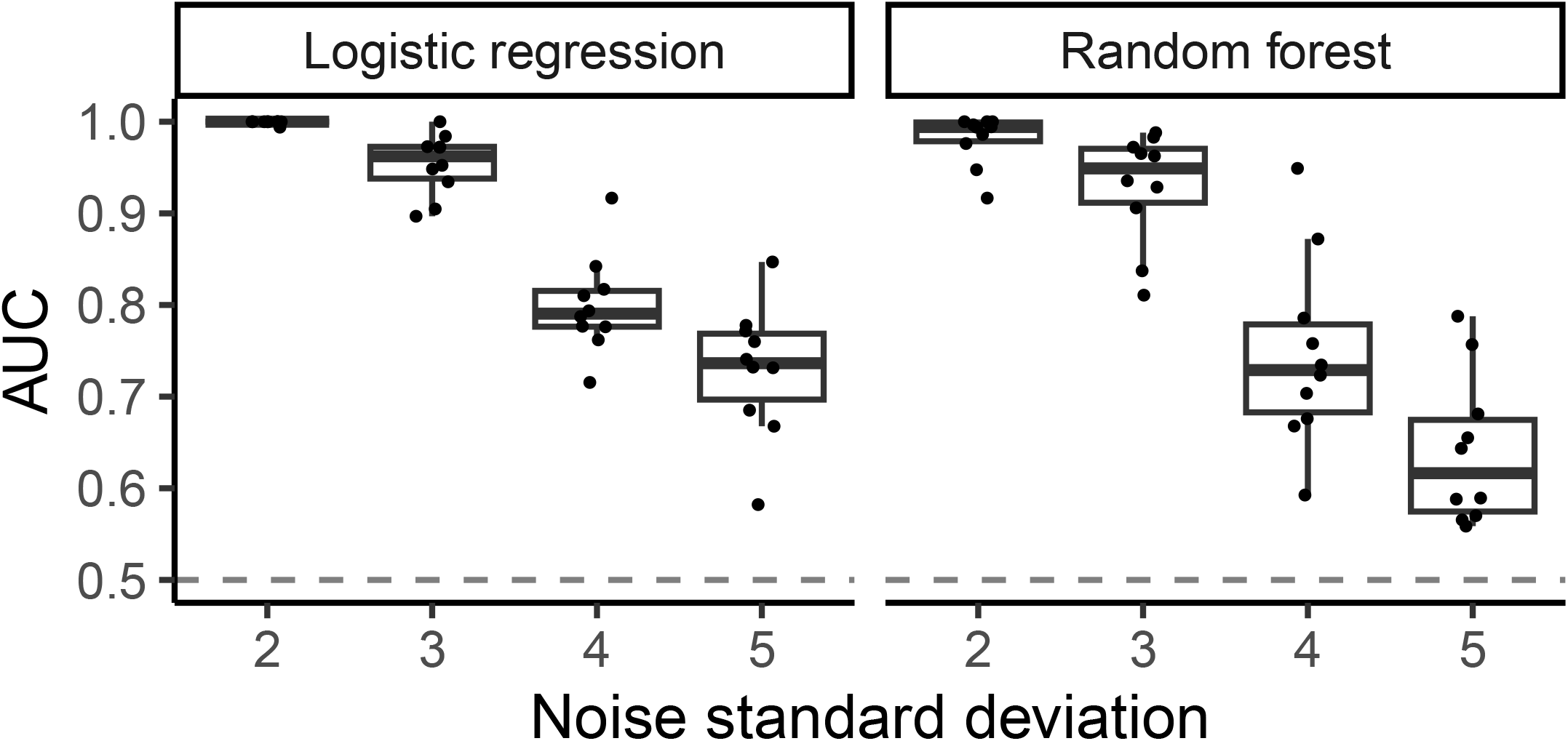
Performance evaluation of `MUSTARD` at different noise levels.

**Figure S2:**
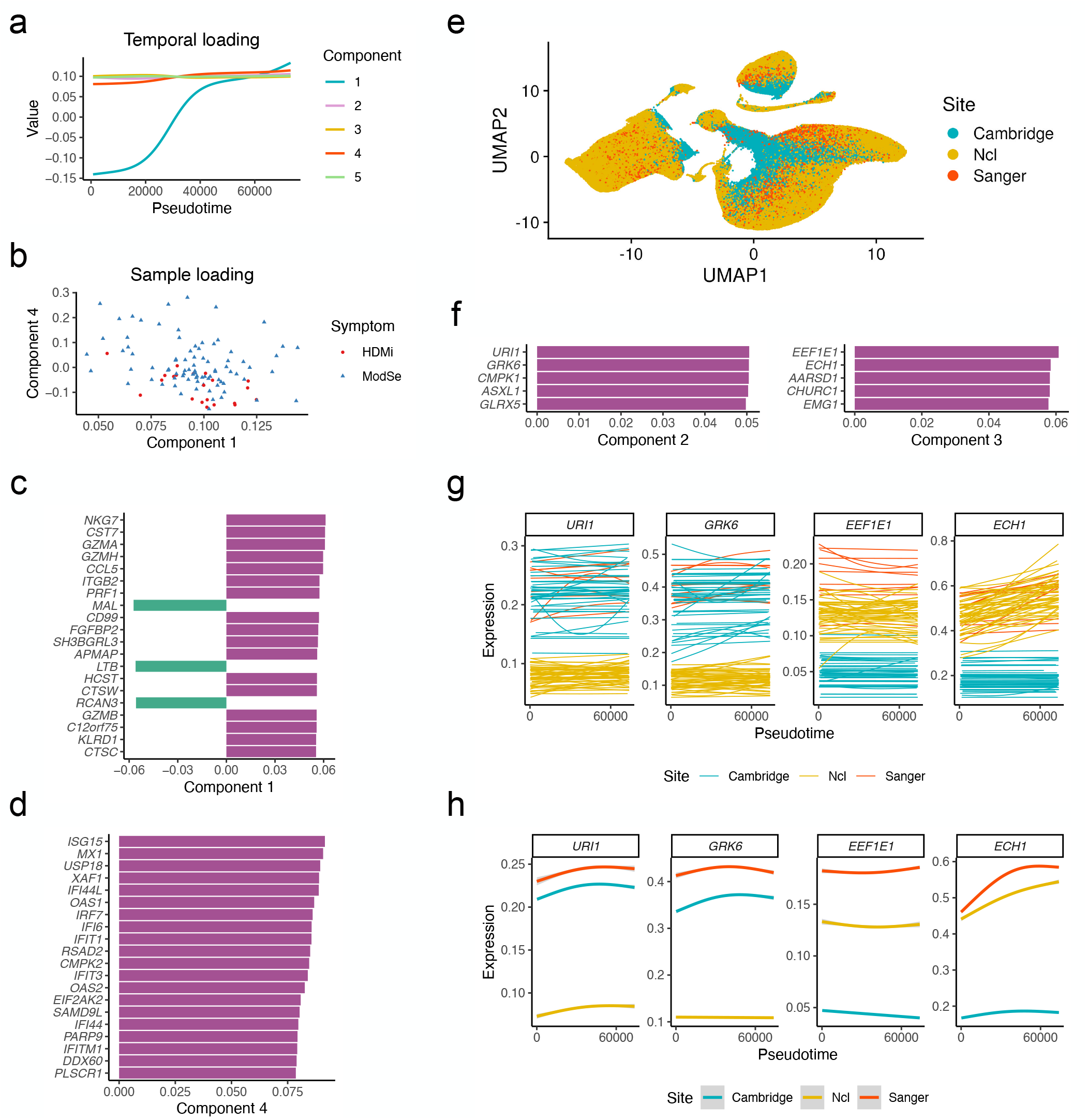
Additional results of COVID-Stephenson study. **a**, Temporal loadings capture major temporal patterns. **b**, Component 4 separates samples from different severity levels. *p*-value obtained by Wilcoxon rank-sum test is 2.14 *×*10^−5^. **c-d**, Top genes in Component 1 (**c**) and Component 4 (**d**). **e**, UMAP showing cells colored by three sites. **f**, Top genes in Component 2 and 3. **g-h**, Example genes’ temporal patterns for each sample (**g**) and each group (**h**).

